# Spatially dispersed synapses yield sharply-tuned place cell responses through dendritic spike initiation

**DOI:** 10.1101/157453

**Authors:** Reshma Basak, Rishikesh Narayanan

**Affiliations:** Cellular Neurophysiology Laboratory, Molecular Biophysics Unit, Indian Institute of Science, Bangalore 560012, India.

**Keywords:** Active dendrites, computational model, dendritic spikes, hippocampus, place cells, synaptic clustering

## Abstract

The literature offers evidence for a critical role of spatially-clustered iso-feature synapses in eliciting dendritic spikes essential for sharp feature selectivity, with apparently contradictory evidence demonstrating spatial dispersion of iso-feature synapses. Here, we reconcile this apparent contradiction by demonstrating that the generation of dendritic spikes, the emergence of an excitatory ramp in somatic voltage responses and sharp tuning of place-cell responses are all attainable even when iso-feature synapses are randomly dispersed across the dendritic arbor. We found this tuning sharpness to be critically reliant on dendritic sodium and transient potassium channels and on *N*-methyl-D-aspartate receptors. Importantly, we demonstrate that synaptic potentiation targeted to afferents from one specific place field is sufficient to effectuate place-field selectivity even when intrinsically disparate neurons received randomly dispersed afferents from multiple place-field locations. These conclusions proffer dispersed localization of iso-feature synapses as a strong candidate for achieving sharp feature selectivity in neurons across sensory-perceptual systems.

Highlights
- Systematic analysis of four synaptic localization profiles for sharp spatial tuning
- Dendritic spikes are generated even with randomly dispersed iso-feature synapses
- Sharp feature tuning could be achieved even in the absence of synaptic clusters
- Targeted, yet dispersed, synaptic plasticity sufficient for place-cell emergence

## eTOC Blurb

The literature exemplifies a critical role of spatially-clustered iso-feature synapses in eliciting dendritic spikes towards sharp feature selectivity, with apparently contradictory evidence demonstrating spatially-dispersed iso-feature synapses. Basak et al reconcile this contradiction by showing that dendritic spike generation and sharp tuning are attainable even when iso-feature synapses are randomly dispersed.

A prominent hypothesis spanning several perceptual systems implicates spatially clustered synapses in the generation of dendritic spikes (dSpikes) that mediate sharply-tuned neuronal responses to input features (DeBello et al., 2014; Druckmann et al., 2014; Govindarajan et al., 2011; Losonczy et al., 2008; Makino and Malinow, 2011; Takahashi et al., 2012; Wilson et al., 2016). On the other hand, there are significant lines of evidence, spanning several perceptual systems, for similarly-tuned synaptic inputs to be dispersed across the dendritic arbor (Chen et al., 2011; Domnisoru and Tank, 2016; Grienberger et al., 2015; Hill et al., 2013; Jia et al., 2010; Varga et al., 2011). How do we reconcile these apparently contradictory observations where spatial clustering is postulated to be required for eliciting dSpikes, and synaptic localization is shown to dispersed? Additionally, a predominant dogma on pyramidal cell neurophysiology is that linear or supralinear modes of operation are respectively preferred when synapses are dispersed or clustered, where the supralinear mode of operation recruits dSpikes owing to cooperativity among concomitantly activated synaptic inputs (Grienberger et al., 2015). Is spatial clustering of synapses an essential component for the expression of concomitantly active synaptic inputs towards the generation of dSpikes? Are spatial clustering of synapses and dSpikes essential in achieving sharp tuning to input features? How do the different voltage-gated ion channels and synaptic receptors that are expressed in the dendritic arbor of pyramidal neurons contribute to sharp tuning to input features?

In this conductance-based morphologically-precise computational study, we tested the hypothesis on the link between spatially clustered synapses and sharply tuned responses by systematically analyzing the impact of distinct synaptic and channel localization profiles on sharpness of spatial tuning in hippocampal pyramidal neurons.

We found that sharply-tuned firing responses were achieved in models where synapses were all clustered on the soma or were randomly dispersed across the dendritic arbor, but not in cases where the same set of synapses were localized to one or two obliques. Strikingly, dSpikes were more prevalent when synapses were randomly dispersed, with sharpness of spatial tuning mediated by the generation and propagation of dSpikes reliant on dendritic sodium channels, transient potassium channels and *N*-methyl-D-aspartate (NMDA) receptors. We confirmed these results with independent multi-parametric stochastic search algorithms spanning thousands of unique models for each synaptic localization scenario. Finally, we also quantitatively demonstrate that synaptic potentiation targeted to afferents from one specific place field to be sufficient to enforce place-field selectivity, even when intrinsically disparate neurons received randomly dispersed afferents from multiple place-field locations.

Our results provide clear lines of quantitative evidence that spatial clustering of synapses is neither essential for the generation of dSpikes nor a requirement for sharp tuning of neuronal responses to input features. We argue that this ability of neurons to achieve nonlinear input processing and sharp feature selectivity with randomly dispersed iso-feature synapses equips them with significant degrees of freedom towards achieving sharp tuning. These quantitative lines of evidence also dispel the impression that dispersed and clustered synaptic localization strategies exclusively translate to linear and nonlinear modes of dendritic operation, respectively. Together, we postulate distinct advantages for the dispersed localization strategy, especially for spatial tuning in the adult hippocampus where new place cells are formed in an experience-dependent manner.

## RESULTS

As a first step in addressing questions on the impact of channel and synaptic localization profiles on sharpness of place-cell tuning profiles, we employed a morphologically realistic conductance-based model of a CA1 pyramidal neuron with channel distributions and physiological measurements (Fig. S1, Table S1) that matched their electrophysiological counterparts (Hoffman et al., 1997; Magee, 1998; Magee and Johnston, 1995; Narayanan and Johnston, 2007; Rathour and Narayanan, 2014; Spruston et al., 1995). We scaled receptor conductances such that the unitary EPSP amplitude at the soma was of 0.2 mV amplitude (Fig. *S1H–I*) irrespective of synaptic location across the dendritic arbor (Andrasfalvy and Magee, 2001). In systematically assessing the impact of synaptic localization profiles on place cell tuning, we placed synapses activated by the same place field at different locations across the somatodendritic arbor and computed sharpness of tuning in the neuronal firing rate for each of these localizations. To quantify the sharpness of the neuronal response, we employed two measurements: the full-width at half maximum (FWHM) and the maximal firing rate (*F*_max_) of the neuron’s firing rate profile (Fig. *1A*).

**Figure 1.**
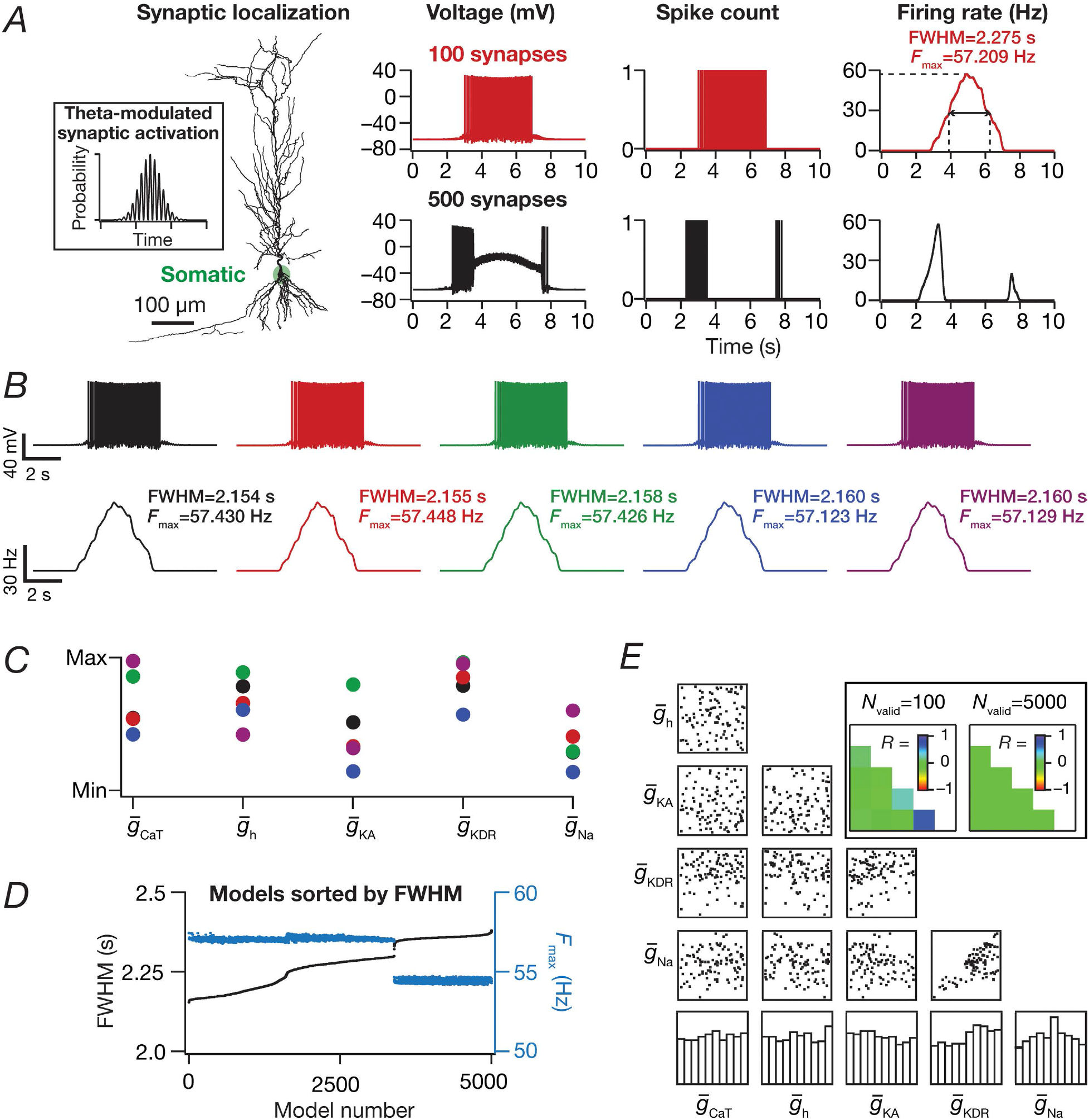
Place field synapses clustered on soma resulted in sharply-tuned place cells with disparate combinations of voltage-gated conductances. (*A*) Left to right: morphological reconstruction of the model with synaptic localization highlighted in green; voltage traces obtained with 100 (*red*) and 500 (black) synapses; spike count plots (1 ms bin); firing rate profiles. (*B*) Five example voltage traces (top) and corresponding firing rate profiles (bottom) of valid models showing similar tuning. (*C*) Normalized parameter values of model cells shown in *B* (same color code). (*D*) FWHM and the corresponding *F*_max_ for all 5000 models, plotted in ascending sequence of their FWHM. (*E*) Scatter plot matrix and matrix representing corresponding Pearson’s correlation coefficients (inset, left) for *N*_valid_=100 models. The lowest row of boxes depicts the distribution of each parameter for all 5000 models. The correlation coefficient matrix plotted for *N*_valid_=5000 models (inset, right) depicts a reduction in correlation coefficients with increase in *N*_valid_ (inset, left). *N*_syn_=100 for *B-E*.

### Models with synapses clustered at the soma elicited sharply-tuned place fields with disparate combinations of channel conductances

As a first step, we placed several (*N*_syn_=100 or 500) conductance-based synapses, receiving afferent presynaptic activity as Gaussian-modulated theta waveform (Eq. 13) from the same place field (Geisler et al., 2010), on the soma (Fig. 1*A*). We found that the base model was capable of eliciting sharp place-cell-like firing responses when *N*_syn_ was 100, but entered depolarization-induced block when *N*_syn_ was 500 (Fig. *1A*).

Was this sharp tuning that was obtained with somatic localization of synapses critically reliant on the specific conductance values set in the hand-tuned base model? Were there explicit constraints on channel conductances to elicit sharply tuned place field responses with somatic localization of synapses? To explore this, we implemented a multi-parametric multi-objective stochastic search (MPMOSS) algorithm (Foster et al., 1993; Goldman et al., 2001; Marder and Taylor, 2011; Mukunda and Narayanan, 2017; Rathour and Narayanan, 2012a, 2014) on all the five active channel conductances 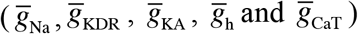 and their gradients (Table S2). We generated 5000 models with each of these maximal conductance values randomly picked from independent uniform distributions spanning 0.5–2 fold of their respective base model values. We activated somatically localized synapses (*N*_syn_=100) with stochastic place-field inputs to each of these 5000 models and calculated FWHM and *F*_max_ of their responses.

A representative set of 5 such model responses indicated similar sharp tuning across all these models (Fig. *1B*). However, the channel conductances that governed these models exhibited wide-ranging variability (Fig. *1C*), implying that sharp place field tuning elicited by somatic localization of place-field synapses was not critically reliant on the specific values of channel conductances. To further confirm this, we plotted the FWHM and *F*_max_ of all 5000 models, and found minimal variability in these measurements suggesting similar tuning (Fig. *1D*). Next, we picked 100 sharply-tuned (low FWHM and high *F*_max_) models and asked if there were pairwise correlations between these model parameters. We found weak pairwise correlation coefficients (max *R*=0.652; min *R*=−0.238; mean ± SEM=0.05 ± 0.08) across all conductance values in these sharply-tuned place cell models (Fig. 1*E*). As all models manifested similar tuning properties, we performed the correlation analysis for all 5000 models, and as expected (Mukunda and Narayanan, 2017; Rathour and Narayanan, 2014), the correlation values grew weaker with increase in the number of models (max *R*=0.007, min *R*=−0.021 with *N*_valid_=5000; mean ± SEM=−0.004 ± 0.002). Together, these results demonstrated that with somatic localization of place-field synapses, disparate channel combinations could yield similar tuning profiles with weak pairwise correlations between the underlying channel conductances.

### Spatially clustered inputs on one or two oblique dendrites did not confer sharpness in place cell tuning

Motivated by lines of evidence suggesting functional dendritic clustering of similar afferent inputs across different neurons (DeBello et al., 2014; Druckmann et al., 2014; Takahashi et al., 2012; Wilson et al., 2016), we placed synapses (*N*_syn_=100 or 500) with identically stochastic (Eq. 13) activation profiles either on a single oblique or split equally across two distinct obliques. Irrespective of whether synapses were placed on one or two obliques, and irrespective of the number of total synapses, we found that the firing rate was low and the place-field profile flat (Fig. *2A*, Fig *2F*), implying weak tuning of the place cell response. Given the critical role of oblique dendritic channel conductances in mediating dSpikes (Losonczy and Magee, 2006; Losonczy et al., 2008), could this conclusion be an artifact of the specific set of channel conductances in the obliques? To explore this possibility, we performed two different MPMOSSs spanning all the conductances 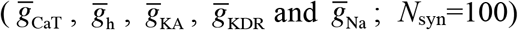 in the one or two obliques, respectively depending on whether synapses were localized onto one (5-parameter MPMOSS; Fig. *2B–E*) or two (10-parameter MPMOSS; Fig. *2G–J*) obliques.

**Figure 2.**
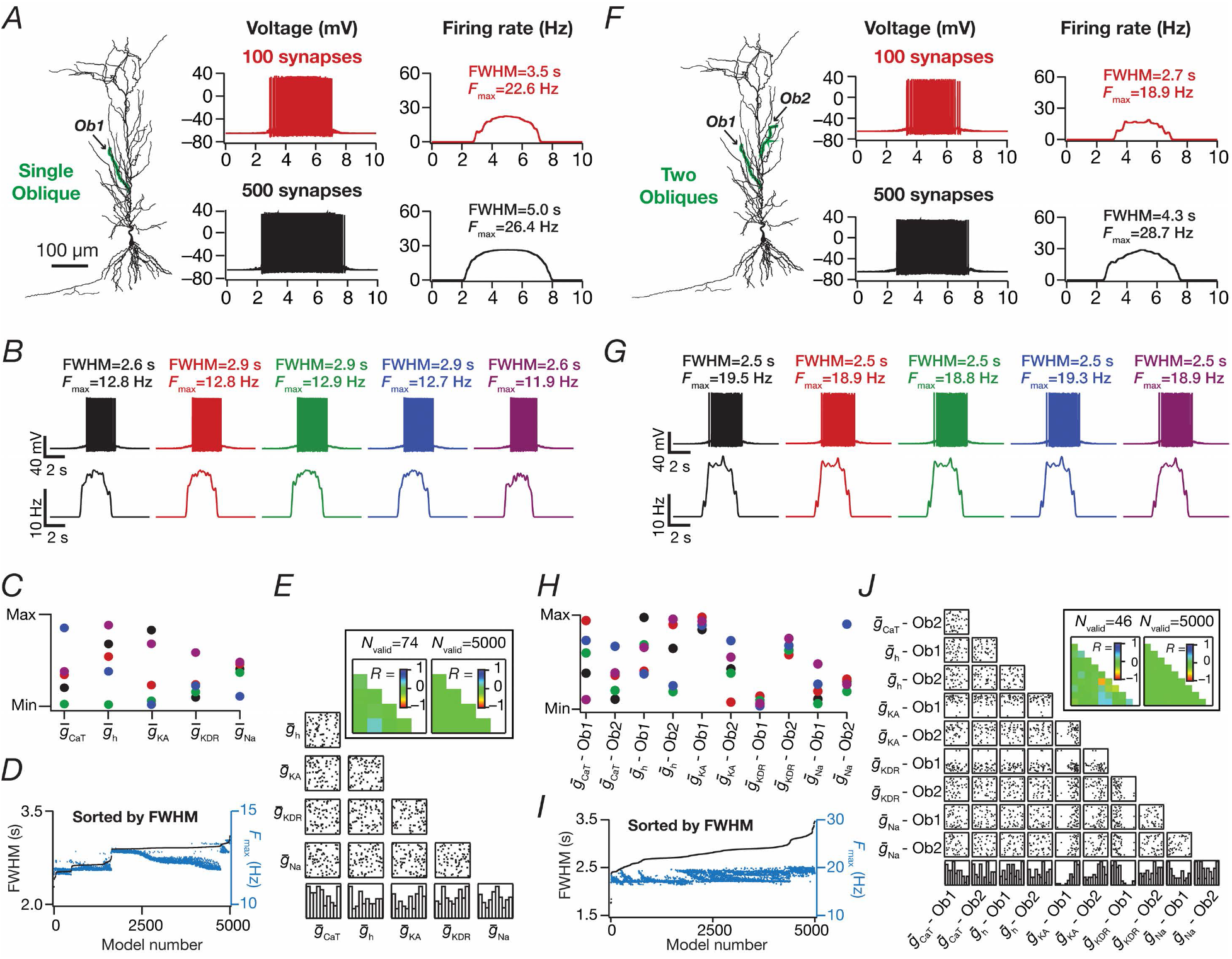
Place field synapses clustered on one or two obliques yielded weak place cell tuning with low firing rates with disparate combinations of local subthreshold channel conductances. (*A*) Left to right: morphological reconstruction of the model with synaptic localization highlighted in green; voltage traces obtained with the different number of synapses; firing rate profiles. *N*_syn_=100 (red), 500 (black). (*B*) Five example voltage traces (top) and corresponding firing rate profiles (bottom) of cell models showing similar tuning profiles. (*C*) Normalized parameter values of model cells shown in *B* (same color code). (*D*) FWHM and corresponding *F*_max_ for all 5000 models, plotted in ascending sequence of their FWHM values. (*E*) Scatter plot matrix and corresponding correlation coefficients (inset, left) for *N*_valid_=74 similarly best-tuned models. The lowest row of boxes depicts the distribution of each parameter for all 5000 models. *N*_syn_=100 for *B-E*. Plots *A-E* correspond to synapses placed on a single apical oblique (*A*, *Ob1* branching from the trunk at ~160 μm from the soma). (*F-I*) Same as *A-E*, but with 50 synapses each clustered on two different obliques (*F*, *Ob1* and *Ob2* branching from the trunk at ~160 μm and ~250 μm, respectively, from the soma) each. Scatter plots and correlation coefficients are for *N*_valid_=46 similarly best-tuned models. In *E* and I, the correlation coefficient matrices plotted for *N*_valid_=5000 models (inset, right) depicts a reduction in correlation coefficients with increase in *N*_valid_ (inset, left).

A representative set of 5 such model responses indicated that the tuning was weak across these models irrespective of whether synapses were localized on one (Fig. *2B*) or two (Fig. 2*G*) obliques although the corresponding channel conductances that governed these models manifested wide-ranging variability (Fig. 2*C*, Fig. 2*H*). This implied that the weak place field tuning elicited by synaptic localization on one or two obliques was not critically reliant on the specific values of channel conductances (Fig. *2D*, Fig. 2*I*). We found weak pairwise correlation coefficients (one oblique: max *R*=0.26; min *R*=−0.1; mean ± SEM=0.009 ± 0.03, *N*_valid_=74; two obliques: max *R*=0.3; min *R*=−0.7; mean ± SEM= −0.02 ± 0.03, *N*_valid_=46) across all conductance values in the similarly best tuned of these place cell models (Fig. 2*E*, Fig. 2*J*). The correlation values grew weaker with increase in the number of models employed to compute the correlation coefficients (one oblique: max *R*=0.007; min *R*=−0.02; mean ± SEM=0.004 ± 0.002, *N*_valid_=5000; two obliques: max *R*=0.03; min *R*=−0.034; mean ± SEM=0.0005 ± 0.002, *N*_valid_=5000). Together, these results provided ample lines of evidence that spatially clustered inputs confined to one or two dendritic branches were incapable of confering sharpness in place cell tuning.

### Dispersed synaptic inputs result in sharply tuned place cell responses

As a next step, we randomly distributed the same set of synapses (*N*_syn_=100) throughout the proximal apical region (radial distance from soma ≤ 300 μm) of the dendritic tree, with all of them receiving afferent place cell activity identically stochastic (Eq. 13) to earlier scenarios. There were 399 possible synaptic locations within this proximal apical region with large intersynaptic distances (Fig. *3B*), which precluded the possibility of incidental clustering of synapses when they were randomly dispersed. We computed the somatic firing rate response and found place cell responses to be sharply tuned (Fig. 3*C*; *cf*. Fig. 1*A*), irrespective of the specific randomization (within the 300 μm location) of dispersed synaptic localization (Fig. 3*C*). When we increased the number of synapses to 200, we noted a reduction in spike height, especially at the center of the place field as a consequence of high excitability and inadequate recovery of sodium channels from inactivation.

**Figure 3.**
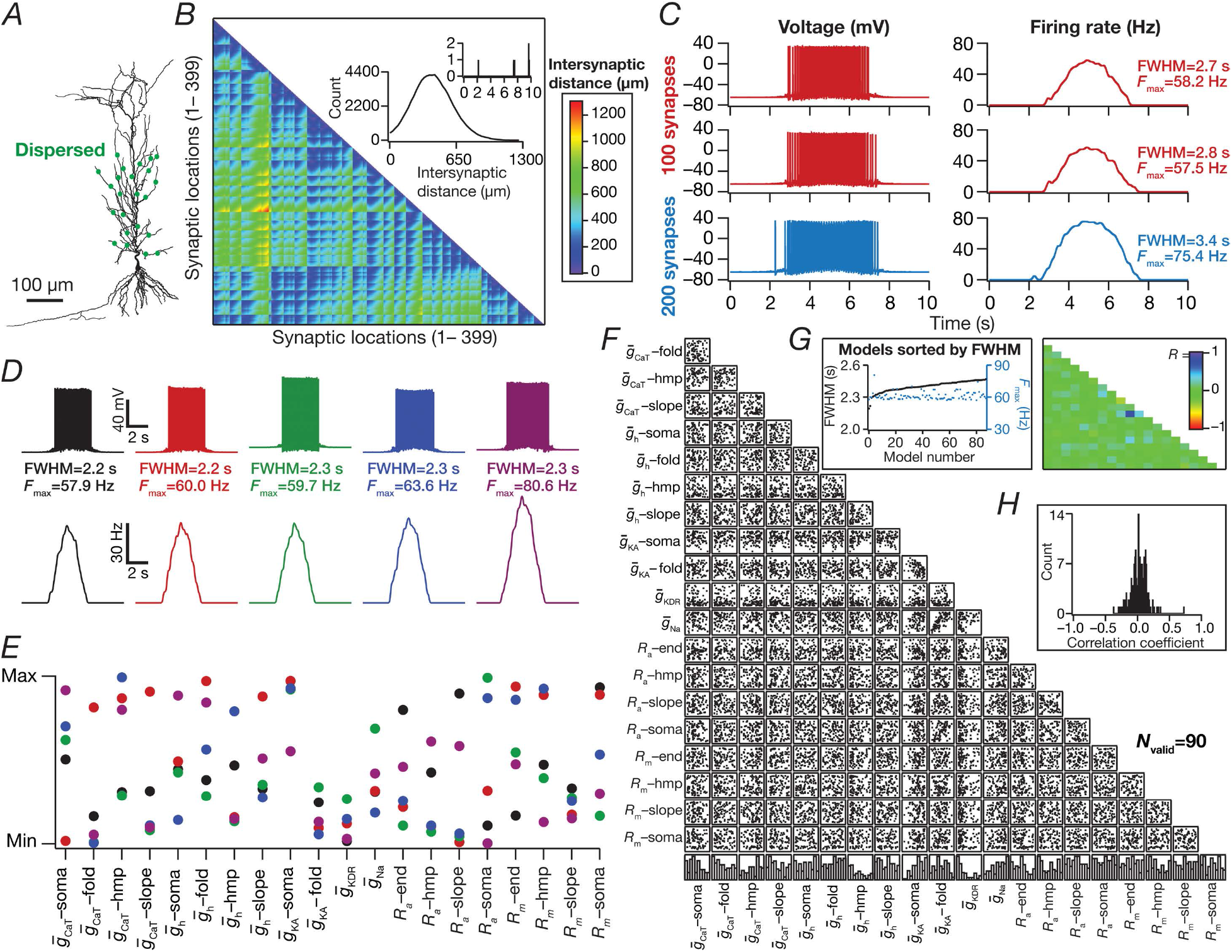
Place field synapses randomly dispersed across the dendritic tree yielded sharply-tuned place cells with disparate combinations of active and passive parameters with weak pair-wise correlations. (*A*) Morphological reconstruction of the model with synapses distributed throughout the proximal 300 μm of the apical dendritic arbor (green dots). (*B*) Lower triangular part of the intersynaptic distance matrix in models where synapses were randomly dispersed across the first 300-μm of the apical dendritic arbor. *Inset*, Histogram of intersynaptic distances plotted in the matrix (top, histogram zoomed to 0-10 μm, showing only 4 intersynaptic distances ≤ 10 μm). (*C*) Left to right: voltage traces obtained with 100 (red) and 200 (blue) synapses; firing rate profiles. The two sets of red traces correspond to two different randomized distributions of 100 synapses. (*D*) Five example voltage traces (top) and corresponding firing rate profiles (bottom) of valid models showing similar tuning profiles. (*E*) Normalized parameter values of model cells shown in *D* (same color code). (*F*) Scatter plot matrix for *N*_valid_=90 similarly best-tuned models, obtained with spatially dispersed place-field synapses (*N*_syn_=100), depicting pair-wise distributions between parameters. The lowest row of boxes depicts the distribution of each parameter for these valid models. (*G*) *Left*, FWHM and corresponding *F*max for the 90 valid models, plotted in ascending sequence of their FWHM values. *Right*, Pearson correlation coefficient (*R*) matrix of the scatter plots in *A*. (*H*) Histogram of the correlation coefficients represented in *G*.

Was this conclusion on sharp tuning with dispersed synaptic inputs an artifact of the specific choice of channel conductances and their localization profiles in the base model? To answer this, we performed an MPMOSS on 20 intrinsic parameters (Table S2) spanning 5000 models with randomly dispersed synaptic localization. We selected 90 of these models as best-tuned valid models based on similarly low FWHM (<2.5 s) and high *F*_max_ (> 55 Hz). We found that despite the similarly sharp tuning profiles of five different cells picked from this population of 90 (Fig. *3D*), the underlying parametric distributions that defined these models covered the entire span of their respective ranges (Fig. 3*E*). Finally, we assessed pairwise correlations of the parameters underlying the 90 valid models (Fig. *3F–H*) and found weak pair-wise correlation across all assessed parameters (max *R*=0.7; min *R*=−0.4, mean ± SEM= 0.01 ± 0.01, *N*_valid_=90). Together, these results provide clear lines of evidence for multiple realizability of sharply tuned place field responses with dispersed synaptic localization profiles, with different randomized distributions of synapses and with disparate channel localization profiles.

### Tuning sharpness and excitatory ramp in somatic voltage response were dependent on the number of synapses

How dependent were our conclusions on the number of synapses? Do these models exhibit an excitatory ramp in their somatic voltages, a characteristic feature in place cell recordings (Bittner et al., 2015; Harvey et al., 2009; Mehta et al., 2000)? Does the somatic voltage trace reflect the theta modulation from the synaptic drive that the neurons receive? Do answers to these questions depend on the specific synaptic localization profile? Would a reduction in number of synapses in the one- and two-oblique localization cases enhance tuning sharpness? We employed valid models from the three distinct MPMOSS procedures (Figs. 2–3) and found that the *F*_max_ and FWHM reduced with reduction in the number of synapses, irrespective of synaptic localization profile (Fig. S2). Importantly, irrespective of number of synapses associated with a place field, *F*_max_ was the largest and the FWHM was the least with the dispersed localization strategy compared to either synaptic clustering strategy.

Next, we smoothed somatic voltage traces of all valid models (from Figs. 2–3), activated with different number of synapses, to eliminate spikes and asked if they exhibited an excitatory ramp (Fig. 4). Although none of the models with any of the localization strategies elicited action potentials (AP) when they were activated with 10 synapses, they exhibited a subthreshold excitatory ramp of a few mV in amplitude (Fig. *4A–B*, Fig. 4*D*). As the number of activated synapses increased, the amplitude of the excitatory ramps expectedly increased, with much larger and sharper ramps achieved with the dispersed localization strategy than with the single-or double-oblique clustering strategy (Fig. 4*B*, Fig. 4*D*).

**Figure 4.**
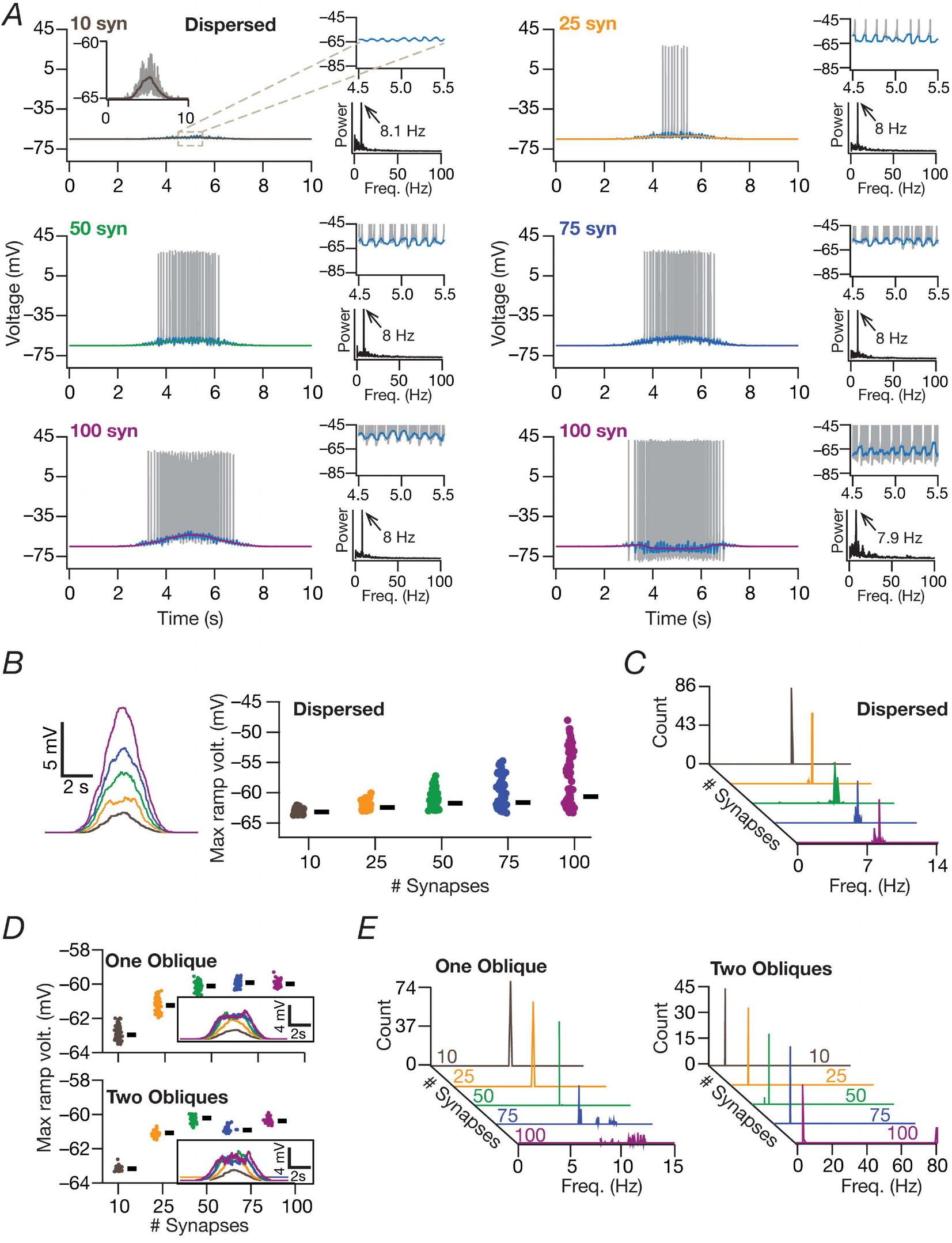
Models with place field randomly dispersed synapses exhibited excitatory voltage ramps, with the ramp amplitude and tuning sharpness dependent on the number of dispersed synapses. (*A*) For all subpanels: *Left*, Excitatory ramp (color-coded traces) overlaid on voltage traces (grey) of a valid cell model activated with different number of synapses. *Right*, top traces depict magnified views of a shorter time window at the place field center, depicting the theta modulation of the ramp (cyan traces). The bottom plots depict the frequency spectrum of the cyan trace (over 10 s time period) also depicting the frequency with maximum power (black arrow). In a small subset of models (100 synapses, right), ramp-like depolarization was suppressed by the dominant afterhyperpolarization dynamics. (*B) Left*, Representative ramps for different numbers of synapses, same as ramp-like traces shown in *A. Right*, Beeswarm plots of maximum ramp voltages for all valid models, plotted for different number of synapses (*N*_valid_=87). Black rectangles depict medians for the corresponding population. (*C*) Histograms of maximum-power frequency in theta-smoothed ramps for all the valid models as a function of number of synapses. For *A-C*, place field synapses were randomly dispersed. (*D-E*) Same as *B-C*, but for models where place field synapses were clustered on one (*N*_valid_=74) or two (*N*_valid_=46) obliques. For *B-E*, color codes for number of synapses are the same as *A*.

Finally, to assess theta modulation of somatic membrane potentials, we smoothed the somatic voltage traces enough to eliminate spikes but to retain theta oscillatory patterns. We performed Fourier analysis on these smoothened traces and found the peak frequency of these spectra (Fig. 4*C*, Fig. *4E*). We found strong ~8-Hz (Eq. 13) power in most models with dispersed synaptic localization irrespective of number of synapses employed (Fig. 4*C*). However, when synapses were clustered within one or two obliques, we found several models where 8 Hz was not the dominant frequency in these smoothed voltage traces, especially when the number of synapses were higher (Fig. 4*E*). These results implied that temporal precision within the theta range was preserved with dispersed synaptic localization, but not when a large number of synapses were confined within one or two obliques. Together, these results demonstrated the remarkable effectiveness of dispersed synaptic localization in achieving sharp firing profile, sharp excitatory ramps and temporal precision in transfer of synaptic inputs, with such effectiveness invariant to the number of activated synapses.

### Dispersed synaptic localization was sufficient to elicit dendritic spikes

What biophysical mechanisms were responsible for the sharp tuning of place cell responses in models with dispersed synaptic localization? Why did synaptic localization confined to one or two obliques not yield sharp tuning? Motivated by the evidence for the role of dSpikes in sharp tuning of place cells (Sheffield and Dombeck, 2015), we simultaneously recorded voltage traces at various points along the somatoapical arbor corresponding to each somatic AP. We performed these recordings for the four different synaptic localization profiles and analyzed these traces specifically for signatures of dSpikes (Fig. 5). Specifically, whereas a backpropagating AP (bAP) would manifest as a dendritic voltage peak *following* the somatic AP peak, a dSpike that potentially participated in AP generation would express as a dendritic voltage peak that *precedes* the somatic AP peak. When synapses were clustered at the soma (Fig. 1), as expected, all dendritic voltages recorded in conjunction with a somatic AP were attenuating bAPs (Fig. *5A*) across all APs from all valid models (Fig. *5E-F*). Thus, with this physiologically unrealistic localization profile with no synapses in the dendrites, it was possible to achieve sharp spatial tuning without dSpikes.

**Figure 5.**
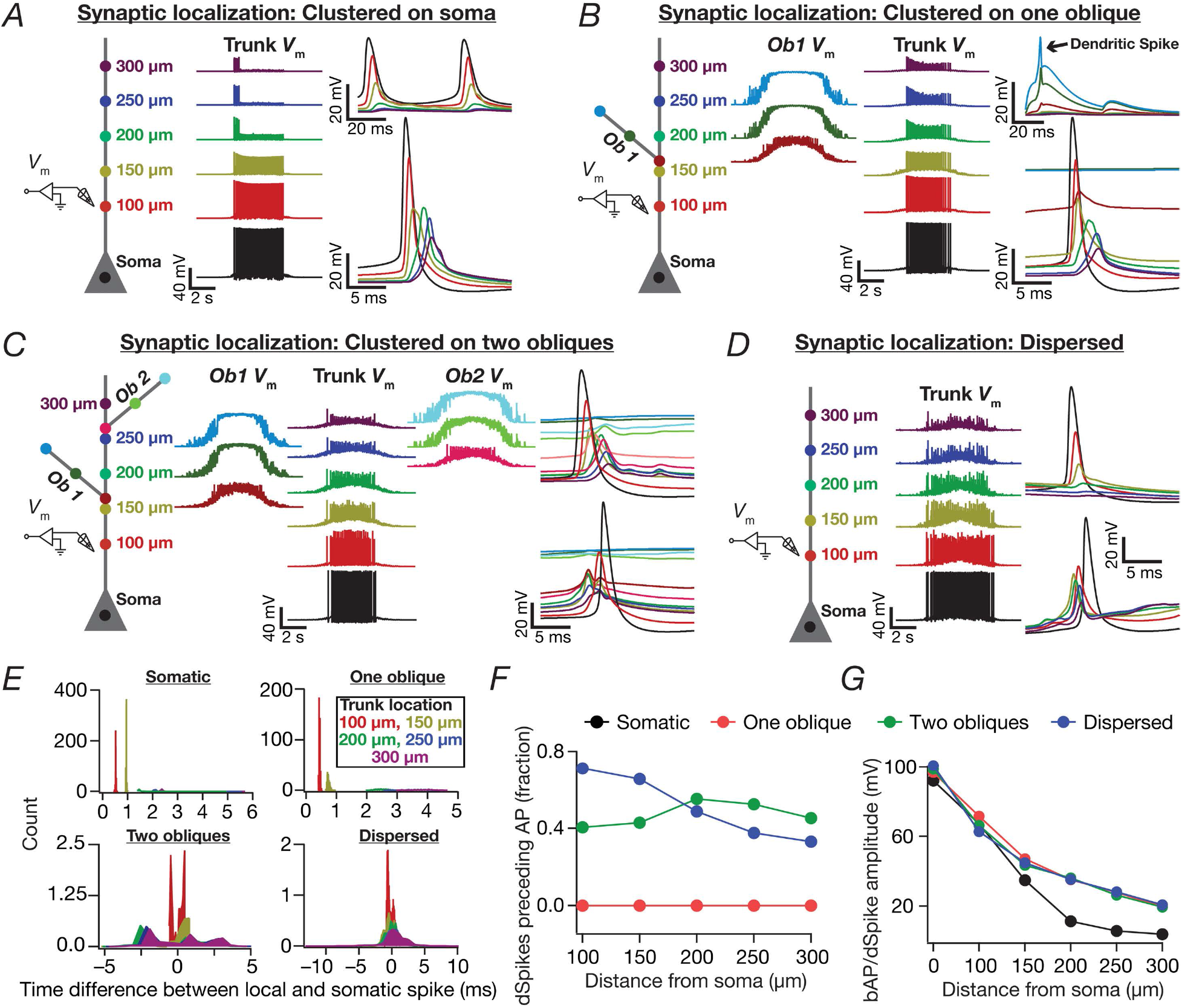
Spatially dispersed synapses yielded sharply-tuned place cell models through dendritic spike initiation. (*A-D*) Color-coded schematic of the neuronal somato-apical arbor (left) showing various points where voltages were recorded (middle and right). *N*_syn_=100 for all cases. *A-D* differ in terms of synaptic localization profiles: (*A*) synapses clustered at soma. (*B*) synapses clustered on one oblique. Shown is a representative non-propagating oblique dSpike (black arrow). (*C*) Synapses clustered at two obliques (50 each). (*D*) Synapses dispersed throughout the somatoapical arbor. For *C-D*, dendritic traces followed (top right) or preceded (bottom right) somatic traces for different somatic APs. (*E*) Histograms of differences between the timing of somatic AP and dendritic bAP/dSpike along the somatoapical axis for the synaptic localization profiles analyzed in *A-D*. Note that negative values indicate the dendritic voltage peak preceding the somatic spike (dSpikes) whereas positive values indicate the somatic AP to precede all other dendritic locations (bAPs). (*F*) Plots, derived from histograms in *E*, depicting the fraction of somatic APs where the dendritically recorded spike preceded the somatic AP. Note that the fraction of preceding dSpikes was zero for all locations when synapses were localized to either the soma or a single oblique. (*G*) Amplitudes of the peak voltages along the somatoapical axis corresponding to the different synaptic localization profiles.

We observed two distinct scenarios when synaptic localization was confined to one oblique, spanning all valid models (Fig. 2*D*). First, during the initial part of the place field when the number of activated synapses was low because of the low probablity of afferent activation, there were dSpikes that were initiated in the oblique where the localization was confined to. However, consistent with *in vitro* evidence (Losonczy and Magee, 2006; Losonczy et al., 2008), these dSpikes did not result in a somatic AP (Fig. 5*B*, Fig. *5E-F*). Second, with progressive increase in activation probability of synapses, the afferent excitation was large enough to drive the oblique to depolarization-induced block. This depolarization traversed through the dendritic arbor to elicit somatic APs, which backpropagated into the dendrites as bAPs (Fig. 5*B*). As somatic firing was elicited by the depolarization, and not by precisely timed dSpikes or synaptic potentials, the firing rate profile was weakly tuned (Fig. 5*B*, Fig. *5E-F*, Fig. 2, Fig. S3). Although the overall scenario was similar to the one-oblique case when synapses were localized on two obliques, there were cases where the dendritic voltage peak preceded the somatic AP peak. These dSpikes were not distinctly observed in the two obliques where the synapses impinged and were observed on the trunk, which then propagated to the soma to elicit APs. However, the large depolarization-induced block introduced in both obliques implied that the tuning was weak when synapses were localized to two obliques (Fig. 5*C*, Fig. *5E-F*, Fig. 2, Fig. S1).

Could the depolarization-induced block observed with single and double oblique localization profiles be because the synaptic drive was large? We repeated our voltage trace analyses with different number of synapses configured to these localization profiles. When the number of synapses was low, the oblique dendrites where synapses were placed showed dSpikes which did not propagate to the cell body (Fig. S3). With increase in number of synapses, the voltage profile within the oblique exhibited depolarization-induced block, and this depolarization travelled to the soma to elicited weakly tuned spatial firing with low temporal precision (Fig. 4*D–E*). Together, functional clustering on one or two oblique dendrites either generated localized dSpikes that did not result in a somatic AP or resulted in a large depolarizing drive that yielded weakly tuned place field responses.

Strikingly, dSpikes preceding somatic APs were more prevalent in the scenario where synapses were dispersed across the dendritic arbor (Fig. *5D-F*). These results demonstrated that widespread depolarization coupled with large-amplitude local EPSPs resulting from the high synaptic strengths (Fig. *S1H-I*) was sufficient to elicit precisely timed dSpikes that together resulted in sharply tuned place field responses.

### Dendritic sodium channels, transient potassium channels and synaptic NMDARs were essential for sharp place cell responses with dispersed synaptic localization

Are apical dendritic sodium channels and NMDARs essential for sharp place cell responses when synapses were dispersed? We employed the virtual knockout model (VKM) technique (Mukunda and Narayanan, 2017; Rathour and Narayanan, 2014) on each of the 87/90 valid models from the MPMOSS algorithm executed with dispersed synapses (Fig. 3). Specifically, we set the apical dendritic NaF conductance or NMDAR permeability in each of these valid models to zero and recorded model voltage and firing responses when identical (to the corresponding valid model) synaptic drive was afferent onto these models. We found that the absence of dendritic NaF channels or synaptic NMDARs resulted in a significant loss of tuning sharpness in these models (Fig. 6*A–G*). As expected, there were no dSpikes when apical dendritic NaF channels were knocked and bAPs did not spread significantly into the dendritic tree. Dendritic spikes were less prevalent in the absence of NMDARs (Fig. 6*A–G*). These results suggest dendritic Na channels and NMDARs as essential ingredients in achieving sharp tuning with dispersed synaptic inputs.

**Figure 6.**
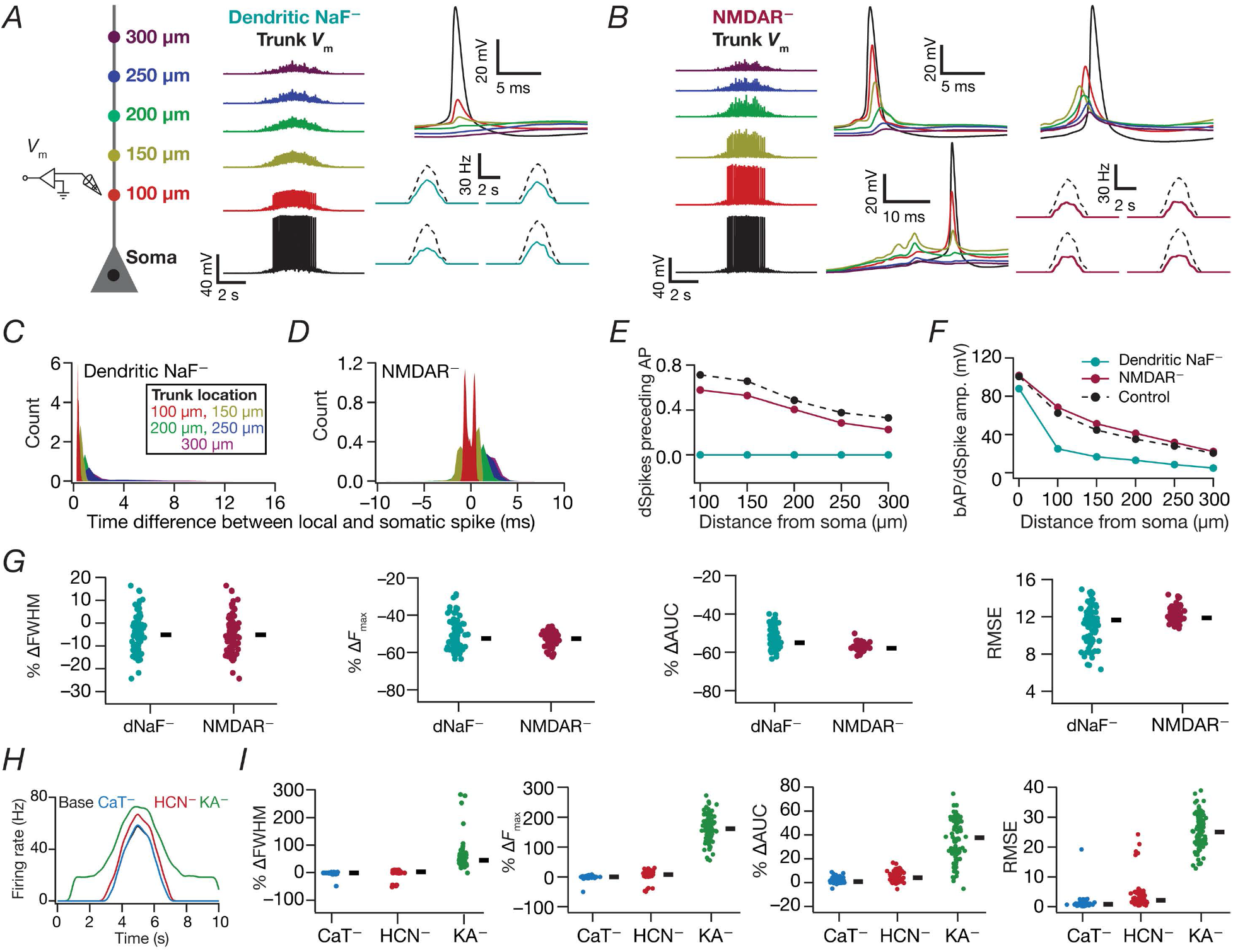
Place cell models with dispersed synaptic localization and disparate channel conductances lost sharpness of spatial tuning in the absence of dendritic sodium channels or NMDA receptors or transient potassium channels. (*A*) Schematic of the neuronal somato-apical arbor (left) showing various points where the voltages (middle) were recorded when apical dendritic sodium (dNaF) channels were knocked out. Firing rate profiles for 4 different model cells (right lower panel) comparing neuronal responses in the presence (black dashed) and the absence (teal) of dNaF channels. (*B*) Voltage traces recorded from various points along the somato-apical arbor (left; color code as in *A*) when NMDA receptors were knocked out across all synapses. Example dendritic voltage traces whose peaks followed (middle top) or preceded (right top) somatic peaks for two different somatic APs are depicted. In some cases dSpikes do not propagate to the soma to generate an AP (middle bottom). Firing rate profiles for 4 different model cells (right lower panel) comparing neuronal responses in the presence (black dashed) and the absence (maroon) of NMDARs. (*C-D*) Histograms of differences between the timing of somatic AP and dendritic bAP/dSpike along the somatoapical axis for the dNaF^-^ and NMDAR^-^ cases. (*E*) Plots derived from the histograms in *C-D* depicting the fraction of somatic APs where the dendritically recorded spike preceded the somatic AP. (*F*) Amplitudes of the peak voltages along the somatoapical axis corresponding to control models and for the dendritic NaF^-^ and NMDAR^-^ cases. (*G*) Beeswarm plots (rectangles are medians) of percentage changes in FWHM, *F*_max_, area under the curve (AUC), root mean square error (RMSE) of the firing rate profiles obtained after virtual knockout of dNaF and NMDARs from each of the 90 valid models in Fig. 3. (*H*) Representative firing rate profile of a valid model (black) and the profiles of the same model cell when different channels were knocked out (color coded). (*I*) Same as *G*, for firing rate profiles obtained after virtual knockout of different channels from each of the 87/90 valid models in Fig. 3.

How sensitive was the sharpness of place cell tuning in valid models with dispersed synaptic localization to each subthreshold ion channel? We employed the VKM technique to measure place cell profiles in each of the 90 valid models by individually setting each subthreshold active conductances 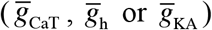 to zero. Although the impact of knocking out these different channels was differential across different models, knocking out the A-type K+ channel yielded the maximum reduction in the sharpness of the place cell tuning (Fig. 6*H*). As expected, knocking out the HCN conductance resulted in a slight increase in excitability, whereas on knocking out the CaT conductance there was a slight decrease in excitability (Fig. 6*H*).

Finally, to check if our results were dependent on the presence of inhibitory synapses inhibition, we incorporated stochastically activated (Eq. 15) GABAAR synapses in the 87/90 valid models obtained with dispersed synaptic localization (Fig. S4). Although we observed an expected reduction in FWHM, *F*_max_, AUC and RMSE (Fig. S4*D*), we noted the prevalence of dSpikes (Fig. S4*C–G*) as well as precise temporal transfer (Fig. S4*C–G*) in the presence of these inhibitory synapses.

### Targeted synaptic plasticity was sufficient to provide selective tuning to a single place field in neurons receiving randomly dispersed inputs from several place fields

If individual place fields projected to independent dendritic branches, clustered plasticity as a mechanism could enable enhancement of synapses only on one of these branches, thereby converting a silent cell (that receives input from several place fields) to a place cell that responds only to one specific location (Bittner et al., 2015; Bittner et al., 2017; Kleindienst et al., 2011; Lee et al., 2012; Losonczy et al., 2008; Makino and Malinow, 2011). However, if the synapses are dispersed as postulated here, then synapses corresponding to a given place field will be distributed all across the dendritic arbor, and would have to undergo simultaneous plasticity. It is feasible to envisage such a scenario because the induction of synaptic potentiation involves *temporal coincidence* of afferent synaptic activation and a postsynaptic depolarization (Magee and Johnston, 1997) without any specific requirement for spatial clustering. Nevertheless, could sharply tuned selectivity to one place field be achieved in neurons that receive afferent inputs from multiple place fields, each of which are randomly dispersed within the dendritic arbor, and are peri-threshold in terms of their ability to elicit place cell responses? Would the overlaps in the spatial locations of synapses from different place fields, and their interactions with the underlying intrinsic somatodendritic properties preclude such selectivity even when synaptic plasticity is targeted only to synapses from one place field?

We independently and randomly dispersed synapses (50 synapses each) from five contiguous place-field locations across the apical dendritic arbor. We had all these place-field inputs to be peri-threshold in the base model, making the model to be a silent cell. With this configuration set as the default base model, we executed an MPMOSS algorithm to generate 2500 models spanning 21 parameters, encompassing the 20 intrinsic parameters (Table S2) and an additional parameter (*permfold*) which governed potentiation of synapses associated with only one of the five (arbitrarily chosen to be the second place field) place field afferents. To validate these models, we accepted models that exhibited selective firing for the second place-field and rejected those that either showed no firing or place-nonspecific firing (Fig. 7*A*). We noted that a significant number (1586 of 2500) of these models were rejected on this criteria of spatially-selective firing, suggesting significant roles for complex interactions of intrinsic neuronal properties with synaptic potentiation and with spatially-dispersed synapses from all place fields. This implies that models that exhibit spatial selectivity are non-trivial demonstrations of how synaptic potentiation could convert a silent cell to a place cell when synapses from several place fields are randomly dispersed only under specific constraints on intrinsic properties. Among models that exhibited spatial selectivity, we then selected those with high *F*_max_ and low FWHM to obtain 60 valid models.

**Figure 7.**
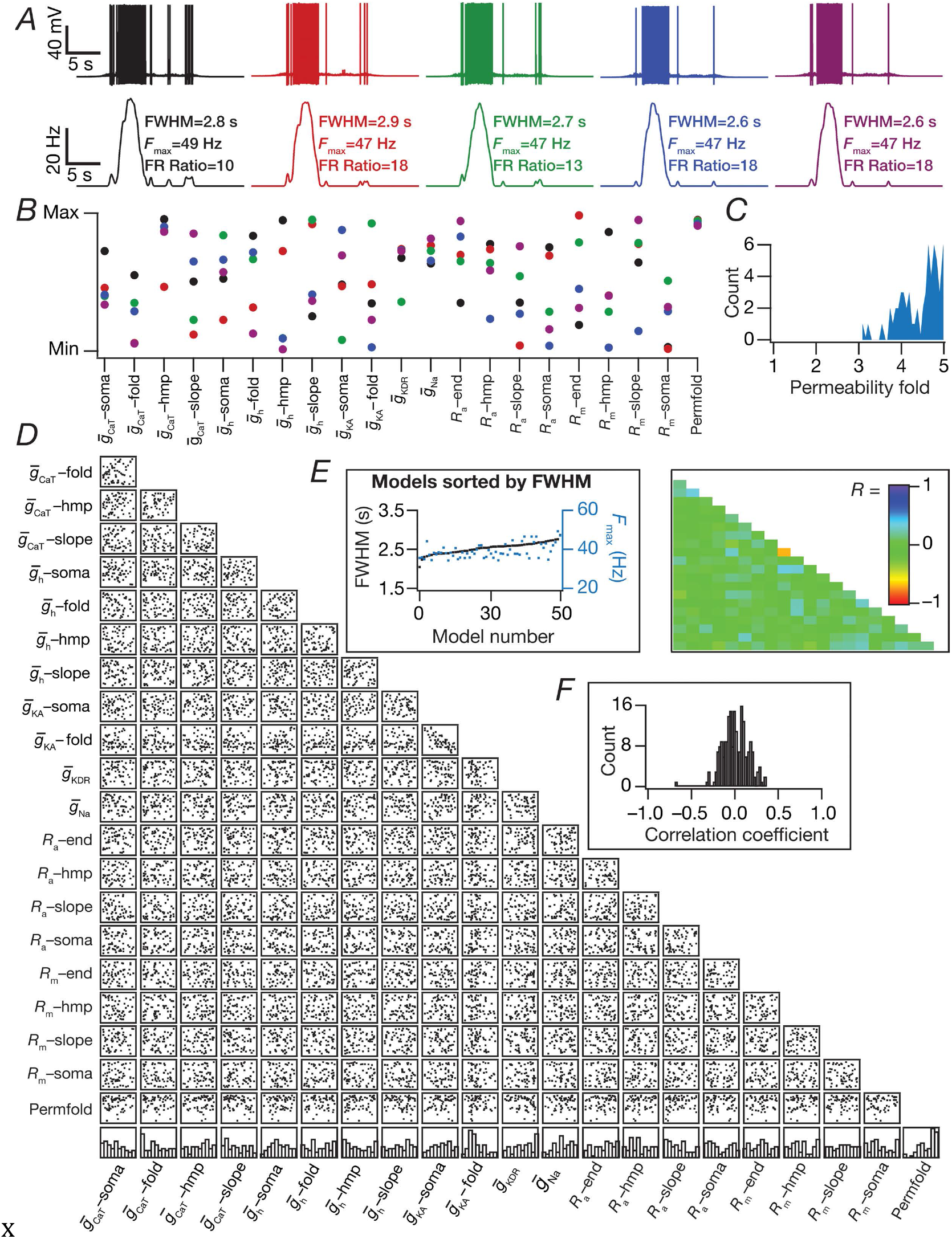
Targeted synaptic plasticity in afferents from a specific place field was sufficient to elicit place-field selectivity in models that received randomly dispersed afferents from multiple place-field locations. (*A*) Five example voltage traces (top) and corresponding firing rate profiles (bottom) of valid models. Note that the firing rate of these model cells within the second place field was at least 10 times larger than their firing rates within any of the other place fields (FR Ratio: ratio between peak in-field and peak out-of-field firing rates). (*B*) Normalized parameter values underlying models shown in *A* (same color code). (*C*) Histogram of the permeability folds (compared to base model permeability) for all the valid models (*N*_valid_=60). (*D-E*) Scatter plot matrix (*D*) and matrix representing corresponding correlation coefficients (*E*) for the valid models. The lowest row of boxes in *D* depicts the distribution of each parameter for all valid models. FWHM and corresponding *F*_max_ for the 60 valid models obtained through MPMOSS, plotted in ascending sequence of their FWHM values (inset in D). (*F*) The histogram of correlation coefficients in *E*.

Are the afore-mentioned constraints on intrinsic properties strong enough to preclude disparate channel combinations from eliciting similarly selective neuronal models? We chose five valid models that exhibited similar sharp tuning to the second place field (Fig. 7*A*), and found significant variability in all model parameters implying a lack of constraints on parameters towards eliciting place-field selectivity and tuning sharpness (Fig. *7B–C*). The valid models also showed significant variability in the level of synaptic potentiation (3–5 fold) required for meeting both validation criteria (Fig. 7*C*). Finally, we observed weak pairwise correlation among the parameters. Together, these results suggest that variable levels of targeted synaptic potentiation, in conjunction with disparate sets of intrinsic properties, was sufficient to provide sharp and selective place-field tuning in a cell that received randomly dispersed inputs from several place fields.

## DISCUSSION

The prime conclusion of this study is that randomly dispersed iso-place-field synaptic localization covering a wide span of dendritic arbor is sufficient to elicit dSpikes and sharply tuned place-field selectivity in neuronal responses. We demonstrated this sufficiency with only a small proportion (*N*_syn_=50−100) of synapses than those that typically impinge (15,000−30,000) on a pyramidal neuron (Bezaire and Soltesz, 2013), with each synapse physiologically constrained (Andrasfalvy and Magee, 2001) in terms of its unitary somatic EPSP. Employing MPMOSS, we demonstrated this sufficiency for different randomizations of dispersed synaptic localization and of disparate channel conductances. The dSpikes and the sharp tuning were critically dependent on dendritic sodium channels synaptic NMDA receptors and A-type K+ channels, providing clear lines of quantitative evidence for a nonlinear mode of neuronal processing despite the dispersion in synaptic localization. Finally, we demonstrated that despite randomly dispersed localization of synapses from several place field afferents, targeted synaptic potentiation was capable of eliciting sharp and selective tuning to one specific place field.

### Synaptic localization strategies and plasticity towards place cell formation

It is evident from our analyses here that the prevalent dogma on the exclusiveness of linear *vs*. nonlinear modes of operation occuring respectively with dispersed *vs*. clustered synaptic localization ignored several features of hippocampal pyramidal neurons. First, the high density AMPARs in dendritic locations implies a large local dendritic voltage deflection corresponding to their afferent activation (Andrasfalvy and Magee, 2001). Second, the opening of slowly-decaying, voltage-dependent NMDARs following this large voltage deflection mediated by AMPARs manifests as a wider spread of a nonlinearly amplified depolarization (Schiller and Schiller, 2001) given that slower signals traverse longer and provide the substrate for better temporal summation (Rall, 1977). Third, given the extents of spatial influence of ion channels and given interactions among depolarizations from multiple dispersed synapses (Rathour and Narayanan, 2012b), the combined large depolarizations emerging from the two receptor subtypes on each synapse was sufficient to cross the threshold for generating dSpikes (Fig. 5). Finally, converging depolarizations and propagating dSpikes from several concomitantly-active dispersed synapses was sufficient to elicit temporally precise somatic APs yielding sharply tuned neuronal responses (Fig. 3–5).

With reference to clustered synapses, it is well established that propagation of single oblique dSpikes, resulting from spatiotemporally clustered synaptic activation, to yield somatic APs is weak, differential and state-dependent (Gasparini and Magee, 2006; Golding and Spruston, 1998; Losonczy and Magee, 2006; Losonczy et al., 2008). The reasoning behind the link between clustered synaptic activation and sharp tuning of neuronal responses is that spatial and temporal summation of multiple such oblique spikes manifests as sharply tuned neuronal firing responses. However, the case for temporal summation of dSpikes from *within* a single oblique is weakened by the presence of dendritic sodium channels whose recovery from inactivation is very slow (Colbert et al., 1997). Such slow recovery from inactivation argues against fast repetitive generation of dSpikes in a single oblique, instead converting large afferent drives into depolarization-induced block (Fig. 2, Fig. 5, Fig. S3). Together, these translate into one of two alternatives when synapses (irrespective of their numbers) were clustered on one or two obliques: a non-propagating oblique spike or a weakly-tuned somatic response consequent to the depolarization traveling from the oblique (Fig. 5, Fig. S3).

In our analyses, we tested synaptic localization profiles confined to one or two oblique dendrites. However, the localization strategies form a continuum from the single oblique cluster to the randomized dispersion ends. Therefore, it is easy to extrapolate the possibility of attaining sharply tuned neural responses with the same number of synapses localized to multiple (not just one or two) obliques (Mel, 1993). This, in turn, points to two distinct somatodendritic synaptic localization strategies towards achieving sharply tuned firing responses through generation and propagation of dSpikes: random dispersion of synapses *vs*. specifically clustered localization limited to a few obliques for each perceptual feature.

Although either strategy would yield sharply tuned responses and it is possible that different neurons employ different strategies, we postulate that there are distinct advantages in choosing the dispersed localization strategy *especially* in the adult hippocampus where new place cells are formed in an activity-dependent manner (Bittner et al., 2015; Bittner et al., 2017; Mehta et al., 1997). First, the rewiring and synapse-formation requirements for targeting afferent synapses corresponding to a newly-forming place field specifically onto a few obliques are more demanding than dispersing these synapses randomly across the dendritic arbor. Second, recent evidence indicates that synapses corresponding to different place fields impinge on silent cells (Bittner et al., 2015; Bittner et al., 2017; Domnisoru and Tank, 2016; Lee et al., 2012). The presence of synapses from several place fields on a single neuron poses a resource allocation problem with reference to specific neuronal surface area assigned to each of these synaptic subsets.

The randomized dispersed localization scenario provides a better solution to the allocation problem rather that assigning multiple of the limited number of obliques to each of these different place fields, especially when the number of place fields to be allocated becomes large. This is possible because of the ability of the neuron to achieve robust tuning with different randomizations of dispersed synaptic profiles and with disparate channel conductances (Fig. 3).

Finally, from a plasticity perspective, the solution involving clusters on multiple obliques seems to offer a distinct advantage given the several demonstrations of clustered synaptic plasticity (Govindarajan et al., 2011; Makino and Malinow, 2011). However, plasticity involves *temporal coincidence* of afferent synaptic activation and a postsynaptic depolarization (Magee and Johnston, 1997), say a widespread plateau potential that even invades the soma (Bittner et al., 2015; Bittner et al., 2017). Importantly, our results offer direct quantitative evidence for the ability of (variable levels of) targeted synaptic potentiation to achieve sharp selectivity to one place field even when multiple place field afferents impinge onto a neuron (with disparate channel properties) in a dispersed manner (Fig. 7). Together, our results offer an advantageous clear alternative to synaptic clustering, with significantly enhanced degrees of freedom, in achieving sharp feature selectivity through dSpike initiation with randomly dispersed iso-feature synaptic afferents.

Although our conclusions are largely extendible to pyramidal neurons in other brain regions, the specifics of morphological characteristics, passive properties, ion channels and receptors expressed there should be rigorously assessed before formulating such an extrapolation. For instance, several cortical pyramidal neurons exhibit characteristic up-down states and the generation of dSpikes through the activation of very few synapses is sufficient to elicit a somatic AP during up states (Palmer et al., 2014). However, hippocampal neurons do not exhibit up-down states, and the cell rests at hyperpolarized resting voltages before the animal enters the place field of the cell that is being recorded (Bittner et al., 2015; Harvey et al., 2009; Lee et al., 2012). As the AP threshold voltage is typically tens of millivolt above threshold, this configuration rules out the possibility of the activation of a few synapses resulting in well-tuned somatic responses in hippocampal neurons. However, given the expression of up-down states in cortical neurons, this should be considered possible in cortical pyramids during up states (Palmer et al., 2014). Future studies should explore the similarities and differences between feature selectivity in different pyramidal neurons across the sensory-perceptual systems with specific focus on the characteristic channels and synaptic properites in different neuronal structure.

## MODELS AND METHODS

Detailed methodological procedures are provided in Supplemental Models and Methods. Briefly, we employed a physiologically and biophysically constrained morphologically precise model of a CA1 pyramidal neuron with synaptic conductances stochastically driven by place-specific theta-modulated presynaptic inputs. We tested four distinct synaptic localization profiles involving iso-place-field synapses: (i) clustered on the soma; (ii) clustered on one oblique dendrite; (iii) clustered on two oblique dendrites; (iv) randomly dispersed across a wide span of the dendritic arbor. We performed separate global sensitivity analyses for each of these four localization strategies, each involving thousands of stochastic models with disparate randomized combinations of channel conductances, and assessed the requirments for sharp tuning of place cell responses. We explored the biophysical mechanisms behind the sharp tuning profile obtained with randomly dispersed synaptic localization profiles by employing virtual knockout models involving dendritic sodium channels, NMDA receptors and each of the dendritically-expressed subthreshold conductances. Finally, we asked if targeted synaptic plasticity is sufficient to elicit sharp and selective tuning to one place field in neurons that received randomly-dispersed inputs from five contiguous place fields.

## CONFLICT OF INTEREST

The authors declare no conflict of interest

## AUTHOR CONTRIBUTIONS

R.B and R. N. designed experiments; R.B. performed experiments and carried out data analysis; R.B. and R. N. co-wrote the paper.

## FUNDING

This work was supported by the Wellcome Trust-DBT India Alliance (Senior fellowship to RN; IA/S/16/2/502727), the Department of Biotechnology (RN), the University Grants Commission (RB) and the Ministry of Human Resource Development (RN).

## ACKNOWLEDGMENTS

The authors thank members of the cellular neurophysiology laboratory for helpful discussions and for comments on a draft of this manuscript.

